# Generative AI for Cell Type-Specific Fluorescence Image Generation of hPSC-derived Cardiac Organoid

**DOI:** 10.1101/2024.01.15.575724

**Authors:** Arun Kumar Reddy Kandula, Tanakit Phamornratanakun, Angello Huerta Gomez, Marcel El-Mokahal, Zhen Ma, Yunhe Feng, Huaxiao Yang

## Abstract

Human pluripotent stem cell (hPSC)-derived cardiac organoid is the most recent three-dimensional tissue structure that mimics the structure and functionality of the human heart and plays a pivotal role in modeling heart development and disease. The hPSC-derived cardiac organoids are commonly characterized by bright-field microscopic imaging for tracking daily organoid differentiation and morphology formation. Although the brightfield microscope provides essential information about hPSC-derived cardiac organoids, such as morphology, size, and general structure, it does not extend our understanding of cardiac organoids on cell type-specific distribution and structure. Then, fluorescence microscopic imaging is required to identify the specific cardiovascular cell types in the hPSC-derived cardiac organoids by fluorescence immunostaining fixed organoid samples or fluorescence reporter imaging of live organoids. Both approaches require extra steps of experiments and techniques and do not provide general information on hPSC-derived cardiac organoids from different batches of differentiation and characterization, which limits the biomedical applications of hPSC-derived cardiac organoids. This research addresses this limitation by proposing a comprehensive workflow for colorizing phase contrast images of cardiac organoids from brightfield microscopic imaging using conditional Generative Adversarial Networks (GANs) to provide cardiovascular cell type-specific information in hPSC-derived cardiac organoids. By infusing these phase contrast images with accurate fluorescence colorization, our approach aims to unlock the hidden wealth of cell type, structure, and further quantifications of fluorescence intensity and area, for better characterizing hPSC-derived cardiac organoids.

## Introduction

Various human organ-specific organoids, including recent cardiac organoids, have been developed and employed in cardiovascular disease modeling and drug screening ^1–5^. To comprehend organoid differentiation and structure, microscopic imaging becomes essential with both phase contrast and fluorescence. In our previous study, a triple reporter human pluripotent stem cells (hPSC) line (3R) was created with three fluorescence reporters for labeling three different types of cardiovascular cells. Upon hPSC differentiation into vascularized cardiac organoids (VCOs), three essential cardiovascular cells can be visualized by live-cell imaging: green (Green Fluorescent Protein - GFP) representing cardiomyocytes (CMs), red/orange (Red Fluorescent Protein/Monomeric Orange Protein - RFP/mOr) representing endothelial cells (ECs), and blue (Cyan Fluorescent Protein - CFP) representing smooth muscle cells (SMCs) ^6^. Since each fluorescence signal corresponds to a specific cell type and corresponding cellular network, this 3R hPSC line has been extremely helpful in tracking cardiac organoid formation in a temporospatial manner for potential applications of disease modeling and drug screening. However, it holds a very obvious limitation, only one cell line is utilized and enabled to visualize the colorful cardiovascular cells in the hPSC-derived cardiac organoids. A diverse and large number of hPSC lines are typically required for achieving more generalized outcomes from biomedical applications. While phase contrast microscopic imaging is routinely applied and conveniently used in every biomedical lab for organoid examination, generating accurate fluorescence information or image colorization on phase contrast images of cardiac organoids would potentially broaden the characterization and analysis of hPSC-derived cardiac organoids in a high-throughput and time-efficient manner.

Numerous approaches have been explored to tackle the challenge of image colorization by artificial intelligence (AI). Traditional machine learning (ML) techniques extract similar features from a reference image to predict colors in a new image ^7–9^ while the efficacy of such methods is contingent on the similarity between the reference image and the target. The advent of Convolutional Neural Networks (CNNs) marked a significant shift, allowing for the automatic extraction of features from images. Pretrained CNNs have gained prominence in image colorization, leveraging feature maps to predict pixel colors ^10–12^. The capabilities of Generative Adversarial Networks (GANs) ^13^ in various generative tasks have prompted their use in colorization. In this context, Conditional GANs, exemplified by Pix2Pix GAN ^14^, have emerged, mapping grayscale inputs to corresponding ground truth images. In our work, we employed Pix2Pix GAN, augmented with the Convolutional Block Attention Module (CBAM) ^15^, enhancing the network’s focus on critical features and elevating colorization realism. Despite the technological advancement of image colorization on generic image categories, there is a lack of research focused specifically on colorizing hPSC-derived tissue constructs, such as cardiac organoids. Small color discrepancies that might be tolerable for generic image generation might be detrimental to cardiac organoids’ images with much smaller features. Even minor color variations in this context can introduce significant misinformation, rendering the task of organoid colorization exceptionally challenging.

Currently, there are three existing techniques for image colorization, including Reference Image-based colorization, which is based on the color information from a reference image. Gupta et al. ^16^ colorized the target images using a reference-colored image, where feature mapping is done for the features extracted using SURF and Gabour filters, and image space voting based on the neighboring pixels is done to obtain the plausible pixel color. This technique suffered at image boundaries and caused color bleeding. To solve this problem, a patch-based feature extraction and colorization technique was proposed in ^17^, that produced more robust colors. With the breakthrough of CNN performance in image processing tasks, CNN is used widely because of its capability to automatically extract the features to find the relations between them and produce more realistic colors. Larsson et al. ^11^ used an architecture from the Visual Geometry Group, VGG-16, in which the first convolution layer was modified to operate on a single channel, and the classification layer was removed. This architecture was fine-tuned on the ImageNet dataset of gray images for one epoch. The grayscale images were passed to this modified network to generate spatially localized multicolumn layers referred to as hypercolumns, which were used to predict the color of the pixel. Multiple CNN architectures, one to extract global-level features of the image and the other to extract mid-level features, created a fusion layer that combined these two global and mid-level features now the final color predictions were generated based on the correlation of those two features ^10^. Moreover, Generative Adversarial Network (GAN) based image colorization is also widely used. Pix2Pix ^14^, a type of Conditional GAN, becomes a popular choice for image colorization because of its ability to find information in pair-to-pair image translation. GAN network consists of a generator, which generates the colorized image from a conditional input image, and a discriminator, which tries to identify if the generated image is real or fake ^18^, GAN architecture was used to generate colored images from infrared images in RGB color space. The pix2pix architecture with U-Net was applied for image colorization on the CIFIAR-10 dataset, where the grayscale image was given as the conditional input to the U-Net generator to generate a colored image, which was then passed to the discriminator to identify if the generated image is a real or fake one ^19^ ^20^.

Accordingly, we established a novel framework utilizing cGANs with adversarial training between the generator and discriminator ^14^ for training on cardiac organoid images (phase contrast and corresponding fluorescence images in green, red, and blue) directly differentiated from 3R, a triple-reporter hPSC line. To address the dynamic nature of cardiac organoid images, we incorporated an attention mechanism, the CBAM ^15^, ensuring an increased emphasis on crucial details and generating more accurate colors. Through an extensive training process on our dataset, the model learned to intricately map grayscale cardiac organoid images (phase contrast) to the corresponding color images (fluorescence). We conducted a thorough evaluation of the proposed method’s effectiveness in preserving biological details and introduced a new evaluation metric, the Weighted Patch Histogram (WPH), designed to capture the color histogram information from small patches of the image, allowing us to obtain a spatially aware color histogram. Collectively, our work demonstrates its efficacy in preserving cell-level information, presenting a promising advancement for the visualization and analysis of hPSC-derived cardiac organoid cell types and structures in biomedical research.

## RESULTS

After hPSC-derived cardiac organoids were differentiated from the 3R triple-reporter hPSC line ^21^, we imaged the entire organoids with live-cell fluorescent microscopy based on three fluorescence reporters: Green (G) – GFP – TNNT2 – CM; Red (R) – mOrange – CDH5 – EC; Blue (B) – CFP – TAGLN – SMC and phase contrast. The 3R hPSC-derived cardiac organoids on day 16 expressed green, red, and blue fluorescence in circular morphology as indicated in **Figure 3**. To increase the diversified results of cardiac organoid differentiation, we included the organoids successfully differentiated into all three cell types (CM, EC, and SMC) and also the organoids composed of two cell types (CM and EC, or CM and SMC, or EC and SMC) and single cell type (CM or EC or SMC). Within this experimental setup, a total of 1,374 paired images (phase contrast and fluorescence) were used for training, while 79 paired images were used for testing and evaluation. To make the model capture the intricate fluorescence details from a limited dataset, the majority of the data was allocated for training. This approach aims to maximize the model’s exposure to diverse examples and the remaining subset of data was used for testing and evaluation to assess the model’s performance.

**Figure 1.**
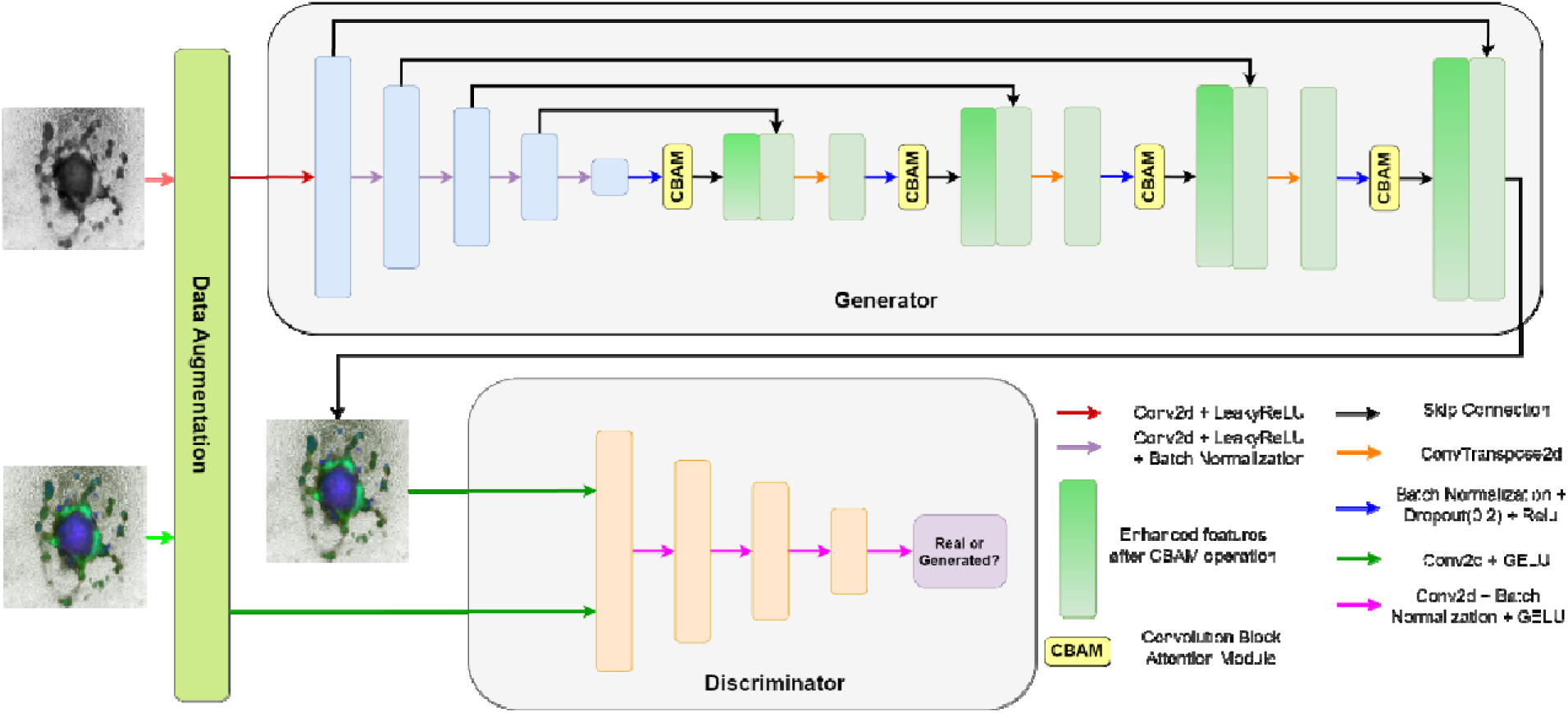
GAN architecture overview of fluorescence colorization of hPSC-derived cardiac organoids.

**Figure 2.**
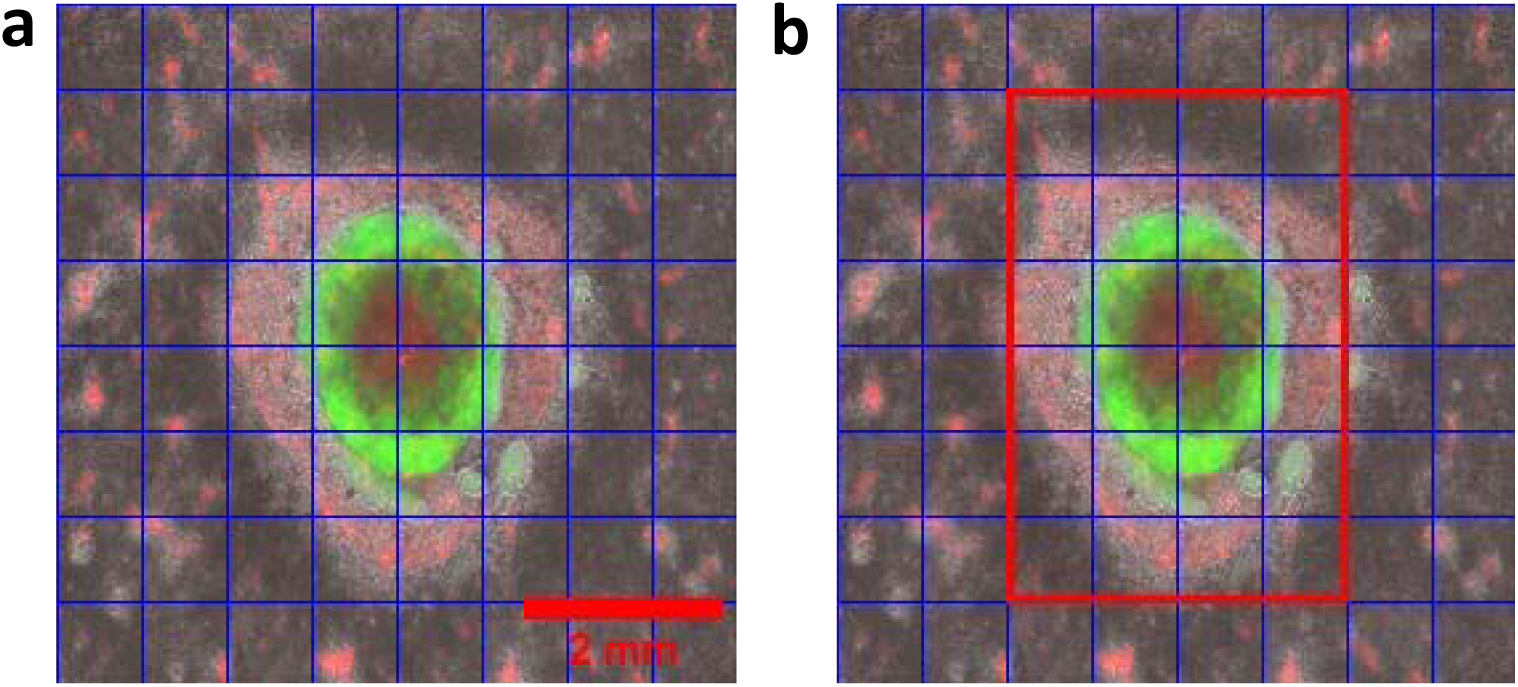
(a) Image divided into 8x8 grid of small patches, (b) Highlighted in red is the ROI, which is given more weightage for histogram comparison. The image size is 256x256, which breaks into a 16x16 grid with multiple patches of size at 32x32 pixels. Scale bar: 2 mm

**Figure 3.**
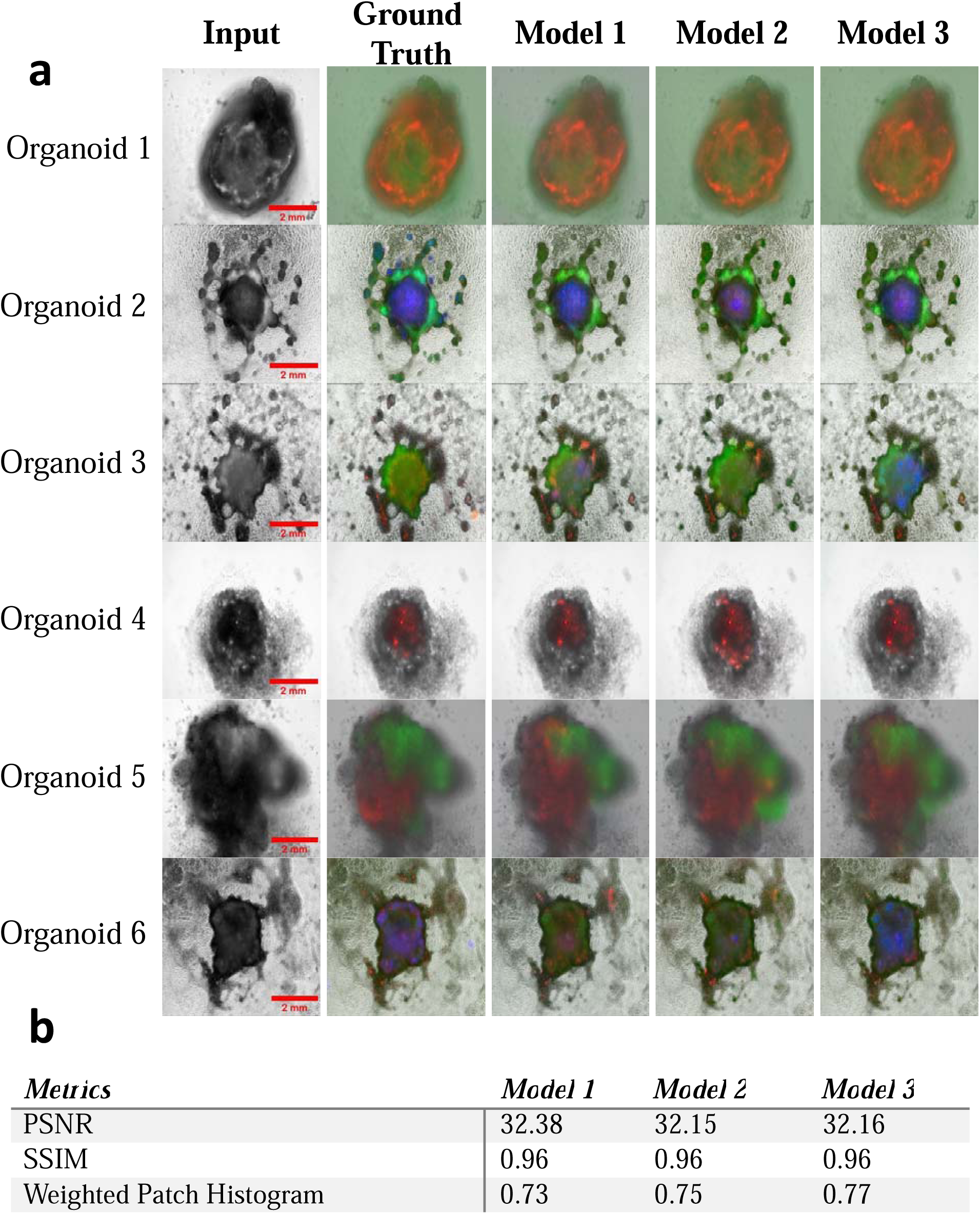
(a) Images of hPSC-derived cardiac organoids from input, ground truth, and predicted images generated by Model 1, Model 2, and Model 3 respectively. hPSC-derived cardiac organoids were not included in the training dataset but from the same batch of experiments. Scale bar: 2 mm (b) Evaluation scores by three different evaluation metrics.

First, we designed three U-Net-based models for optimizing the image colorization of hPSC-derived cardiac organoids:

*Model 1, U-Net generator only:* This generator architecture is based on **Supplementary Fig. 1** without a CBAM block.

*Model 2, U-Net generator with CBAM* as shown in **Supplementary Fig. 1**

*Model 3, U-Net with CBAM and Generator Iteration*. The generator iteration employed the architecture in Model 2 was trained twice in an epoch. With an intuition of making the generator stronger compared to the discriminator as it was trained multiple times to produce more realistic colors.

After three models were trained efficiently with the training dataset of paired phase contrast and fluorescence images from the same organoids, we applied the three models for predicting the hPSC-derived cardiac organoid images in merged phase contrast and fluorescences of green, red, and blue channels as shown in **Fig. 3a** based on the phase contrast images of cardiac organoids. Those organoids used for image prediction and fluorescence colorization were not previously included in the training dataset but from the same batch of organoid differentiation. Three evaluation metrics: the Peak Signal-to-Noise Ratio (PSNR), Structural Similarity Index (SSIM), and WPH were employed to quantitatively evaluate the performance of our models. **Fig. 3b** presents the outcome of evaluation scores achieved on those metrics.

The range of PSNR is [0, ∞], where 0 represents no similarity between images and infinity is for the same images. For a comparison of lossy images, the PSNR score typically ranges between 30 to 50 where the higher the score higher the similarity ^22^. Values over 40 are usually considered to be very good and anything below 20 is unacceptable ^23^. The well-established techniques achieved a PSNR score of 29.52 on the COCO-stuff dataset ^24^, whereas our models achieved PSNR scores are over 32. The COCO-stuff dataset platform is well-known for annotating images or using textual image descriptions by comparing the predicted images to the ground truth of COCO-Stuff at the pixel level. The Structural Similarity Index (SSIM) score ranges in (1, 1) ^25^ where -1 represents no similarity and 1 represents very high similarity. Therefore, a higher score indicates higher similarity. The state-of-the-art techniques have an SSIM score of 0.94 on the coco-stuff dataset ^24^, whereas our models achieved SSIM scores of 0.96. Weighted Patch Histogram ranges in [0,1] where 0 represents no similarity in the histograms of the images, therefore no similarity, and 1 represents full similarity in histograms resulting in a very high similarity of the images. The similarity increases from 0.73 to 0.77 from Model 1 to Model 3.

Since all three models provide good prediction results based on the similarity of the predicted image to the ground truth and evaluation metrics, they were further applied to predict the organoids from different batches of organoid differentiation. As visualized in **Fig. 4a** and quantified in **Fig. 4b**, the predicted organoid images from Model 2 demonstrate the highest similarity in comparison to the other two Models with a higher PSNR (25.26 of Model 2 *vs.* 24.92 of Model 1 *vs.* 24.02 of Model 3) and Weighted Patch Histogram (0.52 of Model 2 vs. 0.49 of Model 1 vs. 0.44 of Model 3). However, the results of evaluation metrics in **Figure 4b** indicate that the scores decrease in terms of all the metrics in comparison to the prediction shown in **Figure 3b**. Since model 2 generated relatively better results than the other two models, model 2 was further fine-tuned by re-training it with one third of images from the new batches of organoid differentiation. After Model 2 was fine tuned, **Fig. 5a** shows the prediction results of the organoids from a new batch of differentiation. It was found that the color generation capability of Model 2 increased after fine-tuning with improved evaluation metrics of PSNR at 29.82, SSIM at 0.94, and WPH at 0.84 (**Fig. 5b**)

**Figure 4.**
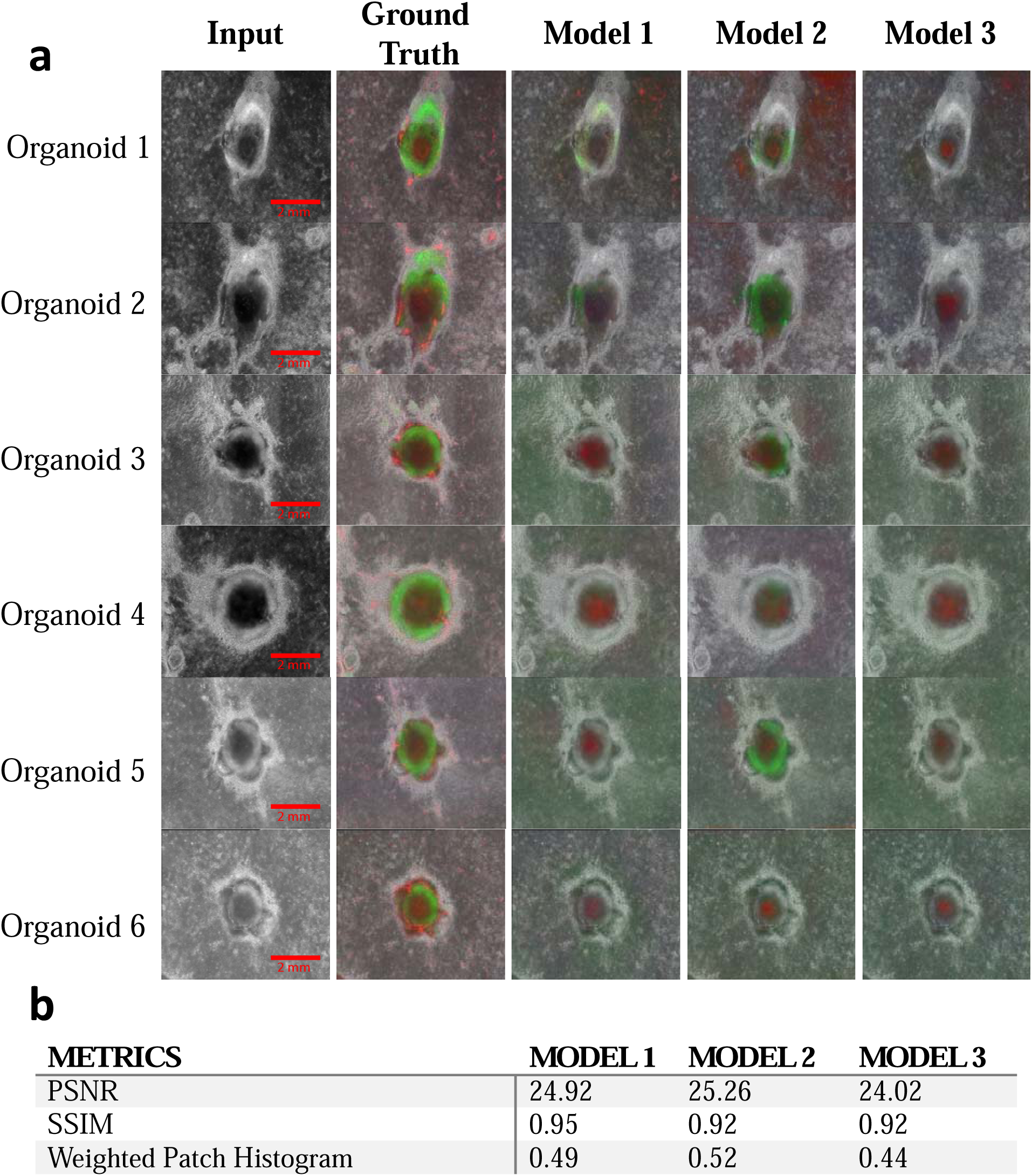
(a) Images of hPSC-derived cardiac organoids from input, ground truth, and predicted images generated by Model 1, Model 2, and Model 3 respectively. hPSC-derived cardiac organoids were from different batches of experiments. Scale bar: 2 mm. (b) Evaluation score on new batches of organoids.

**Figure 5.**
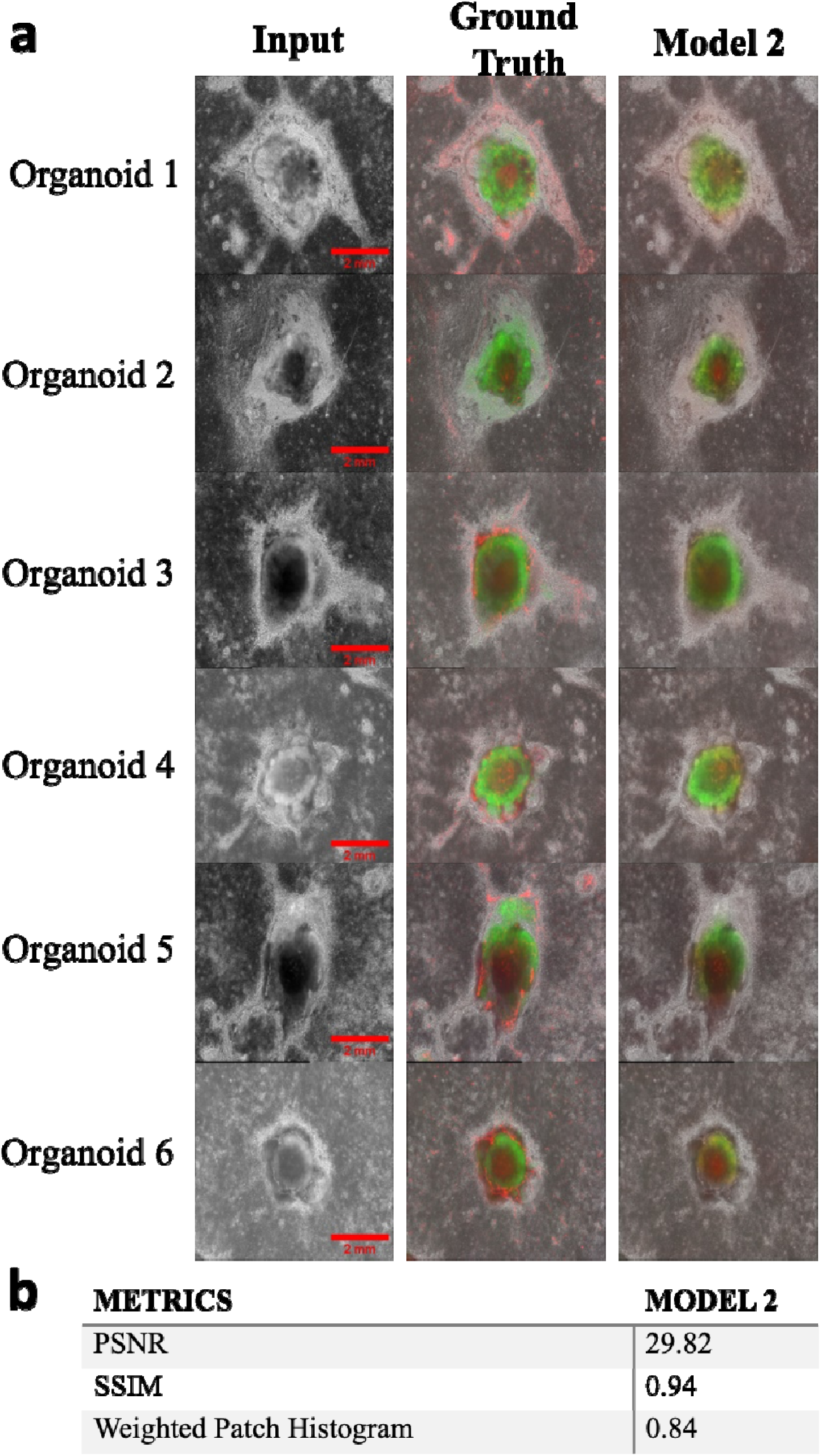
(a) Images of hPSC-derived cardiac organoids after fine tuning from input, ground truth, and Model 2. hPSC-derived cardiac organoids were from different batches of experiments. Scale bar: 2 mm. (b) Evaluation score on new batches of organoids after fine tuning.

To further validate the predicted organoid images by analyzing and quantifying the fluorescence image of each color, which represents one type of cardiovascular cells (CM-green, EC-red, and SMC-blue). The single-channeled fluorescence images were quantified by Organalysis, which is an image processing software for cardiac organoid fluorescence images in high throughput recently developed in our lab ^26^. **Table 1** shows the average results of Organalysis-based analysis ^26^ by comparing the generated images with different metrics like Organoid area, Percentage of Image Covered by Organoid, Total Intensity of Organoid, and Total intensity of Organoid-by-Organoid Area for 70 organoids that were used for the prediction of the same batch of organoid differentiation as shown in **Fig. 3**. The difference% compares the difference between the generated fluorescence and the ground truth of the same organoids, and difference% lower than 25% are highlighted in green blocks and higher than 25% is highlighted in red blocks in **Table 1**. Accordingly, the fluorescence information from the red and green channels generated by Model 1 is close to the ground truth, but not the blue channel. The fluorescence measurements on the red, green, and blue channels generated by Model 2 were close to the Ground truth with difference% in lower or close to 25% in Organoid Area and Percentage of Image Covered by Organoid. Model 3 performed well showing low difference% in all the metrics except for the Total intensity of the Organoid in the Blue channel.

**Table 1.** Quantification and comparison of the individual fluorescence channels in ground truth and predicted images of hPSC-derived cardiac organoids of the same batch of differentiation

|  |  | Organoid Area |  |  | Percentage of Image Covered by Organoid |  |  | Total Intensity of Organoid |  |  | Total Intensity of Organoid by Organoid Area |  |  |
| --- | --- | --- | --- | --- | --- | --- | --- | --- | --- | --- | --- | --- | --- |
|  | Metric | R | G | B | R | G | B | R | G | B | R | G | B |
| Model 1 | Average of Model 1 | 5,287.3 | 12,346.3 | 3,890.1 | 8.1 | 18.8 | 5.9 | 191,929.2 | 647,300.4 | 5,310.3 | 29.8 | 56.9 | 1.9 |
|  | Average Ground truth | 4,813.0 | 12,176.6 | 2,950.6 | 7.3 | 18.6 | 4.5 | 185,273.8 | 674,256.1 | 7,351.5 | 31.3 | 58.0 | 2.9 |
|  | Difference%* | 9.9 | 1.4 | 31.8 | 9.9 | 1.4 | 31.8 | 3.6 | 4.0 | 27.8 | 4.9 | 2.0 | 35.1 |
| Model 2 | Average of Model 2 | 6,057.4 | 14,409.4 | 3,325.4 | 9.2 | 22.0 | 5.1 | 197,167.2 | 704,817.9 | 4,025.1 | 28.4 | 59.6 | 2.5 |
|  | Average Ground truth | 4,813.0 | 12,176.6 | 2,950.6 | 7.3 | 18.6 | 4.5 | 185,273.8 | 674,256.1 | 7,351.5 | 31.3 | 58.0 | 2.9 |
|  | Difference%* | 25.9 | 18.3 | 12.7 | 25.9 | 18.3 | 12.7 | 6.4 | 4.5 | 45.2 | 9.5 | 2.7 | 13.5 |
| Model 3 | Average of Model 3 | 5,615.3 | 13,963.3 | 2,360.0 | 8.6 | 21.3 | 3.6 | 190,031.7 | 675,796.4 | 3,643.2 | 28.5 | 58.7 | 3.0 |
|  | Average Ground truth | 4,813.0 | 12,176.6 | 2,950.6 | 7.3 | 18.6 | 4.5 | 185,273.8 | 674,256.1 | 7,351.5 | 31.3 | 58.0 | 2.9 |
|  | Difference%* | 16.7 | 14.7 | 20.0 | 16.7 | 14.7 | 20.0 | 2.6 | 0.2 | 50.4 | 9.1 | 1.2 | 3.7 |

\*: Green blocks show low difference% less than 25% and red blocks show high difference% more than 25%

Moreover, the cGAN-generated fluorescence information of additional 25 cardiac organoids from a new batch of differentiation in **Fig. 4a** was further extracted and quantified by Organalysis. In **Table 2**, nearly all the models generated the fluorescence information for all three channels at high difference%, which matches with the results of evaluation metrics in **Fig. 4b**. After fine tuning Model 2, **Table 3** shows the results of Organalysis-based analysis by comparing the generated images with different metrics like Organoid area, Total Image Area, Percentage of Image Covered by Organoid, Total Intensity of Organoid, Total intensity of Organoid-by-Organoid Area, and Total Intensity of Organoid by Total Image Area of the organoids in **Fig. 5**. The blue channel for SMC prediction was excluded further due to the inconsistency of the predicted results. More effects in improving the prediction outcome of the blue channel will be achieved in future studies with additional dataset. The image of green fluorescence-labeled CMs in the cardiac organoids generated by the fine-tuned Model 2 is very close to the ground truth with less than a 16% difference to the ground truth in both organoid area and intensity in green fluorescence. The red fluorescence-labeled generated by the GAN model is also close to the ground truth ECs regarding the cardiac organoid area, however, the total intensity of generated red fluorescence is over 30% difference in comparison to the ground truth.

**Table 2.** Quantification and comparison of the individual fluorescence channels in ground truth and predicted images of hPSC-derived cardiac organoids of a new batch of differentiation

|  |  | Organoid Area |  |  | Percentage of Image Covered by Organoid |  |  | Total Intensity of Organoid |  |  | Total Intensity of Organoid by Organoid Area |  |  |
| --- | --- | --- | --- | --- | --- | --- | --- | --- | --- | --- | --- | --- | --- |
|  | Metric | R | G | B | R | G | B | R | G | B | R | G | B |
| Model 1 | Average of Model 1 | 3,909.8 | 25,471.5 | 3,024.1 | 6.0 | 38.9 | 4.6 | 29,111.6 | 381,391.7 | 3,029.6 | 14.7 | 20.3 | 0.9 |
|  | Average Ground truth : | 5,164.9 | 3,258.8 | 300.7 | 7.9 | 5.0 | 0.5 | 145,953.5 | 212,736.2 | 305.8 | 29.3 | 65.4 | 0.8 |
|  | Difference %* | 24.3 | 681.6 | 905.8 | 24.3 | 681.6 | 905.8 | 80.1 | 79.3 | 890.6 | 49.8 | 68.9 | 9.0 |
| Model 2 | Average of Model 2 | 4,658.3 | 12,125.7 | 10,130.4 | 7.1 | 18.5 | 15.5 | 50,595.1 | 232,752.4 | 10,623.4 | 12.1 | 37.5 | 0.8 |
|  | Average Ground truth | 5,164.9 | 3,258.8 | 300.7 | 7.9 | 5.0 | 0.5 | 145,953.5 | 212,736.2 | 305.8 | 29.3 | 65.4 | 0.8 |
|  | Difference %* | 9.8% | 272.1 | 3269.2 | 9.8 | 272.1 | 3269.2 | 65.3 | 9.4 | 3373.5 | 58.8 | 42.7 | 4.1 |
| Model 3 | Average of Model 3 | 1,333.9 | 39,878.6 | 6,444.3 | 2.0 | 60.8 | 9.8 | 22,187.4 | 570,512.7 | 7,885.9 | 16.4 | 13.8 | 0.8 |
|  | Average Ground truth | 5,164.9 | 3,258.8 | 300.7 | 7.9 | 5.0 | 0.5 | 145,953.5 | 212,736.2 | 305.8 | 29.3 | 65.4 | 0.8 |
|  | Difference %* | 74.2 | 1123.7 | 2043.2 | 74.2 | 1123.7 | 2043.2 | 84.8 | 168.2 | 2478.4 | 44.0 | 78.9 | 1.3 |

\*: Green blocks show low difference% less than 25% and red blocks show high difference% more than 25%

**Table 3.** Summary of the quantification and comparison of the individual fluorescence channels in the ground truth and predicted images of hPSC-derived cardiac organoids on a new batch of organoid differentiation

|  | Organoid Area |  | Percentage of Image Covered by Organoid |  | Total Intensity of Organoid |  | Total Intensity of Organoid by Organoid Area |  |
| --- | --- | --- | --- | --- | --- | --- | --- | --- |
| **Metric** | **R** | **G** | **R** | **G** | **R** | **G** | **R** | **G** |
| Average of Model 2 | 4,232.5 | 2,946.3 | 6.5 | 4.5 | 56,174.0 | 121,931.8 | 15.6 | 40.4 |
| Average Ground truth | 4,606.7 | 3,178.5 | 7.0 | 4.9 | 103,742.0 | 166,253.0 | 22.2 | 52.9 |
| Difference%* | 8.1% | 7.3% | 8.1% | 7.3% | 45.9% | 26.7% | 29.7% | 23.7% |

\*: Green blocks show low difference% less than 25% and red blocks show high difference% more than 25%

## DISCUSSION

hPSC-derived cardiac organoids are the most emerging *in vitro* human heart model, which has been used from basic developmental biology to translational drug discovery and regenerative medicine, however, how to characterize hPSC-derived cardiac organoids in high efficiency and efficacy at examining cardiovascular cell type-specific expression and networks without additional fluorescence immunostaining and imaging has not achieved yet. This study filled this gap by introducing a novel strategy for fluorescently colorizing cardiac organoids from phase contrast images by utilizing cGANs and CBAM. The findings of the study illustrate the efficiency of this framework in capturing fluorescence intricacies of the cardiovascular cells (CMs, ECs, and SMCs) in the hPSC-derived cardiac organoids.

To better evaluate the prediction outcomes from the algorithms of cGANs+ CBAM, three different evaluation metrics were applied with varied emphasis and focus on image recognition and comparison. For example, the WPH was included as a new metric to highlight the efficacy of our approach in preserving biological details compared to traditional metrics like PSNR and SSIM. Typically, the images generated with evaluation scores of PSNR over 30, SSIM over 0.92, and a WPH score over 0.75 are the most accurate and similar to the ground truth.

Initially, the prediction of fluorescence images within the same batch of organoid differentiation was highly accurate, especially by integrating the CBAM into the conditional GAN framework of Model 2, which captured salient features in phase contrast cardiac organoid images with significant improvement. This attention mechanism enhances the quality and fidelity of the generated colorizations by directing the model’s focus toward critical regions within the image and generating realistic and accurate colorizations of grayscale organoid images ^27^. To further test the prediction outcome of organoids differentiated from different batches, we included additional organoids from the other two new batches of organoid differentiation. However, the prediction accuracy was greatly reduced in PSNR and WPH. To address this problem and bolster the prediction accuracy of organoids from the different batches of differentiation, we did fine tuning by incorporating one third of organoid images from the new batches of differentiation into the training dataset. This step of fine turning did improve the prediction outcome with higher evaluation metrics.

Finally, to meet the important need for organoid characterization in image quantification, we conducted the fluorescence image analysis and comparison between prediction and ground truth. We adapted the most common measurement of organoid images focusing on cardiovascular-specific cell types: Organoid area, Percentage of Image Covered by Organoid, Total Intensity of Organoid, and Total intensity of Organoid-by-Organoid Area. The percentage of differences (difference%) in Organoid area, Percentage of Image Covered by Organoid, Total Intensity of Organoid, and Total intensity of Organoid-by-Organoid Area are all lower than 25% in the prediction of the same batches of organoids in G (GFP-CMs) and R (mOrange- ECs), however, the difference% of B (CFP-SMCs) in Total Intensity of Organoid is larger than 25% due to the insufficient dataset containing blue fluorescence information in hPSC-derived cardiac organoids. Moreover, the intensity of blue fluorescence was significantly lower than the other fluorescence channels making it highly sensitive in the quantification of blue fluorescence. Similar to the results of evaluation metrics, the difference% in the organoid characterization measurement is more than 25% prior to the fine tuning. Through the optimization of fine tuning, the difference% in green and red fluorescences becomes lower than 10% in the Organoid area and Percentage of Image Covered by Organoid with significant improvement in the fluorescence colorization of hPSC-derived cardiac organoids, however, the prediction of fluorescence intensity-related measurements needs further improvements due to the variation of microscopic imaging at different days and batches even using the same imaging setup and parameters.

## LIMITATIONS AND FUTURE WORKS

While the established cGAN+CBAM algorithm has achieved satisfied predictions of hPSC-derived cardiac organoid fluorescence images from the corresponding phase contrast images, a few limitations still need to be further addressed to improve the prediction accuracy with additional functions. For example, the prediction of blue-SMC fluorescence is still insufficient in both image visualization and quantification, and the cellular network prediction of green-CM and red-EC fluorescence could be further improved as well. To overcome this limitation, we will increase the dataset size with more images at varied sample categories, such as including the cardiac organoids with varied and defined ratios of each fluorescence through controlled organoid differentiation. Also, we will consider employing ensemble learning techniques, where multiple models are trained, and their predictions are combined to improve overall accuracy and robustness. As supported by the results of fine tuning, the prediction accuracy was enhanced significantly, however, how to achieve a promising prediction outcome without fine turning has not been achieved yet. We will try incorporating the Progressive GAN ^28^ technique in our training approach to enhance training stability and capture intricate details of hPSC-derived cardiac organoids to skip the step of fine tuning and still achieve high accuracy of fluorescence colorization. Accordingly, the predicted image quantification related to fluorescence intensity measurement will be improved further for the organoids from a new batch of differentiation. Another limitation of the current study is only epi-fluorescence images were included in the training dataset. In consideration of the three-dimensional (3D) structure of hPSC-derived cardiac organoids, the confocal fluorescence microscopic imaging with a 3D image stack will be considered to predict the 3D structure of organoids with cell-type specific expressions and networks. Lastly, the prediction of cardiac organoids differentiated from more hPSC lines will be included and evaluated to extend the application of this technology.

## CONCLUSIONS

In conclusion, a novel model was established to address the critical challenge of colorizing phase images of hPSC-derived cardiac organoids using cGANs and CBAM. This framework has demonstrated its efficacy in capturing intricate multichannel fluorescence information within the hPSC-derived cardiac organoids, enhancing the interpretability and analysis of cardiovascular cell type and biomarker expression in both images and quantification for biomedical research and applications. The cGAN model, enriched by the CBAM module, outperformed the other two models, showcasing its adaptability and effectiveness by evaluating and comparing three evaluation metrics. Notably, for optimal results on the organoid from new batches of differentiation, fine tuning the model is suggested, ensuring that accurate and faithful fluorescence information is generated. Moreover, the quantification of fluorescence information in predicted organoid images brings extensive validation of hPSC-derived cardiac organoids for broader and impactful biomedical applications, such as the prediction of cell type-specific drug cardiotoxicity, prediction of cardiovascular development, sex, race, and genetic/mutation-specific disease evaluations, if more diverse hPSC cell lines are included in the training dataset. A similar algorithm or strategy can also be applied to the brain, liver, kidney, and cancer organoids for automatic fluorescence colorization and quantification.

## METHOD and MATERIALS

### A. Framework Overview

In this section, we presented an overview of our research on image colorization of phase contrast or grayscale images of hPSC-derived cardiac organoids using cGAN, specifically the Pix2Pix model ^14^.

Our methodology was built on the utilization of the Pix2Pix conditional GAN ^14^, the Pix2Pix model, short for “Pixel-to-Pixel Translation,” is a notable example of a cGAN. To achieve this, we adopt the CIELAB color space, consisting of three channels: Lightness, a*, and b*. In CIELAB, Lightness represents the grayscale channel, while a* and b* represent the two-color channels. This Lightness channel will serve as the conditional input to the generator, and the a* and b* channels will be the target channels for generating colorized versions of the grayscale images. The objective of using CIELAB color space is to extract only the color information from the cardiac organoid and train the model to generate the plausible colors of a* & b* that will be merged on the grayscale input, to obtain the colorized cardiac organoid.

Additionally, we incorporated the CBAM ^15^ to increase the channel and spatial attention of the GAN model to focus on the relevant features. CBAM is an innovative enhancement introduced to the architecture of deep neural networks, particularly CNNs. CBAM integrates both channel and spatial attention mechanisms, facilitating the model’s ability to focus on pertinent features within the input data. Channel attention enables the network to adaptively assign importance to different channels, emphasizing relevant information while suppressing noise. Simultaneously, spatial attention ensures that the network allocates its focus to meaningful spatial regions within an image.

The primary motivation for incorporating conditional GANs, CIELAB color space, and CBAM is to increase the model’s attention to relevant features and limit the model’s predictions to only two channels (i.e a* & b*), thereby reducing the number of predictions compared to the RGB color space, where the model would have to make predictions for the R, G, B channels. The synergy between Pix2Pix, CIELAB, and CBAM contributes to notable colorization outcomes. **Figure 1** illustrates the main components and steps of the process of the image colorization workflow, which depicts the transformation of a grayscale cardiac organoid image to a fully colorized output using Pix2Pix conditional GAN. The conditional input passed to the Generator is the Lightness channel and the Discriminator was trained on the a* & b* channels.

### A. Individual Models

#### U-Net generator

The U-Net generator consists of an encoder and a decoder, connected by a bottleneck layer. **Supplementary Figure 1** demonstrates the architecture of our U-Net generator where the encoder progressively reduces the spatial dimensions of the input grayscale image while extracting features. The decoder then upsamples these features to produce the final colorized output. Skip connections between corresponding encoder and decoder layers facilitate the flow of low-level features, enhancing the network’s ability to capture fine details.

One distinctive feature of the U-Net generator here is its utilization of the Lightness (L) channel from the CIELAB color space as a conditional input. This L channel represents the grayscale information of the input image. By incorporating this channel, the generator can focus on producing color information (a* and b* channels) that is coherent with the grayscale content.

#### Convolutional Block Attention Module (CBAM)

The Generator’s ability was enhanced using the CBAM, which integrates channel and spatial attention mechanisms, enabling the discriminator to adaptively assign importance to different channels and meaningful spatial regions within the image. CBAM as shown in **Supplementary Figure 1** is an integral component incorporated into our U-Net Generator architecture to enhance its ability to capture and emphasize relevant features within grayscale organoid images. The input feature map is F ϵ R^CXHXW^ and CBAM extracts 1D channel attention map M_C_ ϵ R^C X 1 X 1^ and a 2D spatial attention map M_S_ ϵ R^1XHXW^. In summary, the overall attention processes can be explained as:

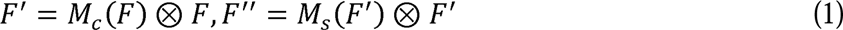

where ⊗ denotes the element-wise multiplication and the resulting F” is the final refined output map that includes the details from both channel attention and spatial attention. This operation allows the model to focus on relevant features while suppressing irrelevant information ^15^.

Channel attention enables the network to adaptively assign importance to different channels of feature maps, emphasizing relevant information while suppressing noise. Channel attention is essential when dealing with multi-channel images such as the L*a*b* color space we operate in. This selective channel weighting allows the model to focus on the most informative colorization components.

Spatial attention is another crucial aspect of CBAM. It ensures that the network allocates its focus to meaningful spatial regions within an image. In the context of colorization, this is especially important as it guides the model to concentrate on the relevant regions where colorization details are essential. Spatial attention complements channel attention by pinpointing critical areas in the input.

#### Patch Discriminator

The Patch Discriminator is a CNN designed to operate on image patches rather than entire images as shown in **Supplementary Figure 2**. This approach allows the discriminator to focus on local details and textures, making it well-suited for assessing the quality of colorizations at a fine-grained level. It consists of multiple convolutional layers to produce a single feature map that is used to classify the patch as real or fake. The final classification result for the entire image is obtained by averaging the predictions from the patches across the entire image. The result is a global classification score that represents the discriminator’s assessment of the overall image

The Patch Discriminator engages in adversarial training with the U-Net generator. It aims to distinguish between real colorized organoid patches and fake patches generated by the generator. Through this adversarial process, the discriminator provides feedback to the generator, encouraging it to produce colorizations that are indistinguishable from real color images.

The primary objective of the Patch Discriminator is to guide the U-Net generator in generating high-quality colorizations. Assessing the local realism of colorized patches helps ensure that fine-grained details and textures are faithfully preserved in the output.

#### Loss Functions

Discriminator Loss: The Discriminator, a key component of our conditional GAN, serves the crucial role of assessing the authenticity of colorized organoid images. To fulfill this role, the Binary Cross-Entropy Loss (BCEWithLogitsLoss) was used.

Mathematically, the discriminator loss can be expressed as:

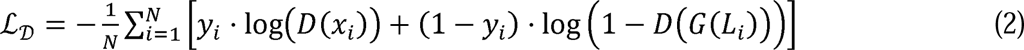

Here, l_D_ represents the discriminator loss, where N is the batch size, x_i_ denotes the ground truth colorized organoid images, i_i_ represents labels for real images i_i_ 1 and fake images ( i_i_ 0 , D x_i_ ) signifies the discriminator’s output for real images, and c L_i_ signifies the generator’s output for the corresponding grayscale input L_i_ . The BCEWithLogitsLoss computes the binary cross-entropy loss by comparing the discriminator’s predictions with the ground truth labels.

The discriminator aims to maximize this loss, which encourages it to correctly classify real and fake patches within the images. Simultaneously, the generator minimizes this loss during adversarial training to produce colorizations that are indistinguishable from real images.

Generator Loss: The Generator, a pivotal component of our conditional GAN, is tasked with generating plausible colorizations. To achieve this, a combination of two loss functions: Binary Cross-Entropy Loss (BCEWithLogitsLoss) and L1 Loss (Mean Absolute Error), were used. Similar to the discriminator, BCEWithLogitsLoss as its adversarial loss function was used. It encourages the generator to produce colorizations that convincingly fool the discriminator into classifying them as real.

Mathematically, the generator’s adversarial loss is defined as:

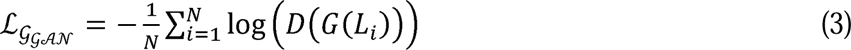

This loss drives the generator to produce colorizations that are perceptually similar to real color images.

L1 Loss (Mean Absolute Error): In addition to the adversarial loss, the L1 Loss was incorporated to ensure that the generated colorizations closely match the ground truth images in terms of pixel-wise similarity.

Mathematically, the generator’s L1 loss is expressed as:

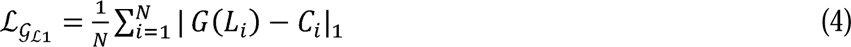

Here, L_GLl_ represents the generator’s L1 loss, where N is the batch size, L_i_ denotes the grayscale input images, c L_i_ represents the generator’s colorized output, and C_i_ denotes the corresponding ground truth color images. The L1 loss encourages the generator to produce colorizations that closely match the ground truth, focusing on fine-grained pixel-level details.

By combining these two loss components, the generator was trained to produce colorized organoid images that are both visually convincing and pixel-wise accurate, ultimately enhancing the quality and realism of the generated colorizations

### B. Image Similarity Measurement Metrics

Evaluating the accuracy and quality of the generated image is a challenging task and on top of that, we have a limited dataset of 1300 images so we used non-deep-learning metrics to obtain a similarity score. We applied 3 different evaluation metrics: PSNR, SSIM, and WPH to compare the similarity between ground truth and colorized images.

PSNR was used to measure the quality of reconstructed or compressed images and this metric is used for comparing the similarity of the colorized image with ground truth ^24,29–32^. It objectively measures how well a colorization technique preserves the details and visual fidelity of the original image. By calculating the PSNR value, we can evaluate the accuracy and fidelity of colorization algorithms. Its range is (0, ∞), 0 represents no similarity between images, and the higher the score higher the similarity.

PSNR score of an *m* x *n* (width x height) image I and its compressed image K can be determined by:

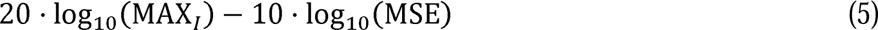

Where MAx_/_ is the maximum possible pixel value of the image and MSE is the mean square error of the Original Image I and its compressed image K, it can be calculated by:

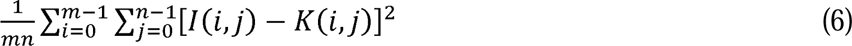

The SSIM is a widely used evaluation metric for assessing the visual quality of the colorized image with ground truth ^24,29–32^. It considers global and local image characteristics, capturing the perceptual differences and structural similarities between the colorized and ground truth images. To be specific, SSIM compares three components in an image pair, suppose x and y are the two patches of the true and compressed image respectively that are aligned with each other, the luminescence comparison function l(x,y) captures the differences in brightness, the contrast comparison function c(x,y) accesses variation in image contrast and the structure comparison function s(x,y) measures differences in image structure and texture. SSIM is a combination of all these three factors ^33^.

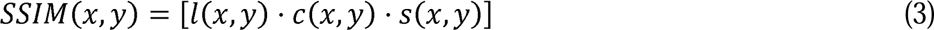

By evaluating the preservation of underlying structures and textures, SSIM provides a comprehensive measure of the algorithm’s ability to maintain visual coherence and realism. The SSIM score typically falls within the range of (-1, 1) ^25^, where a higher score signifies greater similarity.

PSNR and SSIM are widely used metrics in evaluating Image Colorization tasks. Still, they are not exactly appropriate for the problem, because PSNR is designed to identify the quality of the compressed image with the original image. Similarly, SSIM primarily focuses on structural similarity rather than color which is the main part of Image colorization. So, we tested the WPH to compare the similarity of generated colors. With regular Histogram comparison, valuable spatial information of the color is lost, so in our approach, we split the image into a 16x16 grid (**Figure 2(a)**) to have multiple small patches of the image and compare these small patches individually to the corresponding patch from the ground truth. This patch histogram comparison increased the spatial information of the pixel’s value.

As patch histogram comparison increases spatial color information, therefore, reducing the patch size to the smallest possible value may produce the best results. The smallest size possible to compare is 1x1 pixels, which leads us to a pixel-to-pixel comparison of the images and it would be highly sensitive to noise and unreliable. So, the optimized balance between the patch size and the number of bins in the histogram comparison was tested and validated, and a patch size of 32x32 pixels and 32 bins was found ideal for histogram comparison in cardiac organoid images. Since most of the cardiac organoids were centered in the image, and were the region of interest (ROI), which provides enhanced significance in color comparison without the excessive background. The weightage for the patches inside the ROI in **Figure 2(b)** was increased by 50% to give more importance to the colors in the organoids.

### A. Comparison and quantification of prediction image in each cell type

According to our recently published organoid image pre-processing and analysis platform – Organalysis ^26^, the Organoid area, Percentage of Image Covered by Organoid, Total Intensity of Organoid, and Total intensity of Organoid-by-Organoid Area were quantified from the predicted images and paired ground truth of each organoid with the following measurements:

Organoid area: total pixel numbers of fluorescence per cell type in each organoid

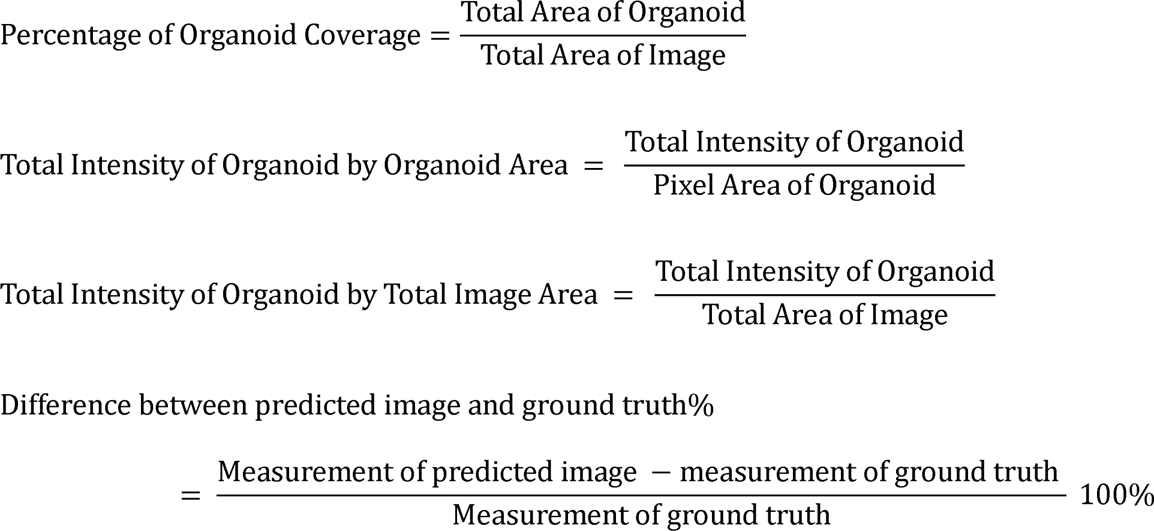

## Source Code

For complete access, please reach out to Yunhe Feng, and Huaxiao Yang

## Contribution Statement

HXY and YHF: Conceptualization; AKRD, YHF, and HXY: writing – original draft; AKRD and TP: formal algorithm establishment and analysis; ZM: manuscript review and editing; RKB, AHG, and MEM: organoid differentiation and characterization.

## Funding Supports

This work was supported by the NIH R15HD108720 (HXY), Harry Moss Heart Foundation (HXY), start-up from the University of North Texas (UNT) Biomedical Engineering (HXY), Research Seed Grants (2021 and 2023) from UNT Research and Innovation Office (HXY), NIH G-RISE T32GM136501 (AHG), NIH R01HD101130 (ZM), and NSF 1943798 (ZM).

## Conflicts of Interest

None

## Supplementary materials

**Supplementary Figure 1.**
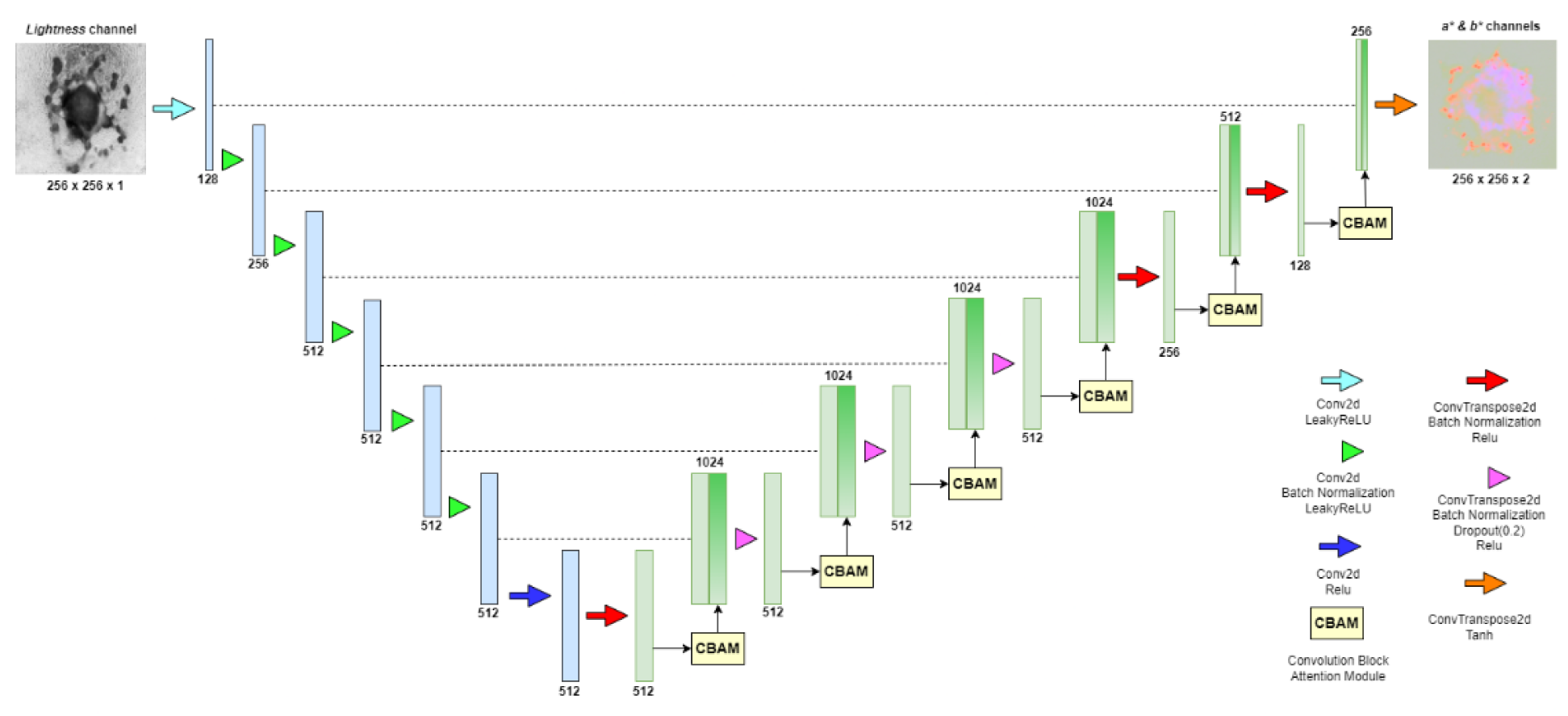
U-Net generator Architecture with CBAM

**Supplementary Figure 2.**
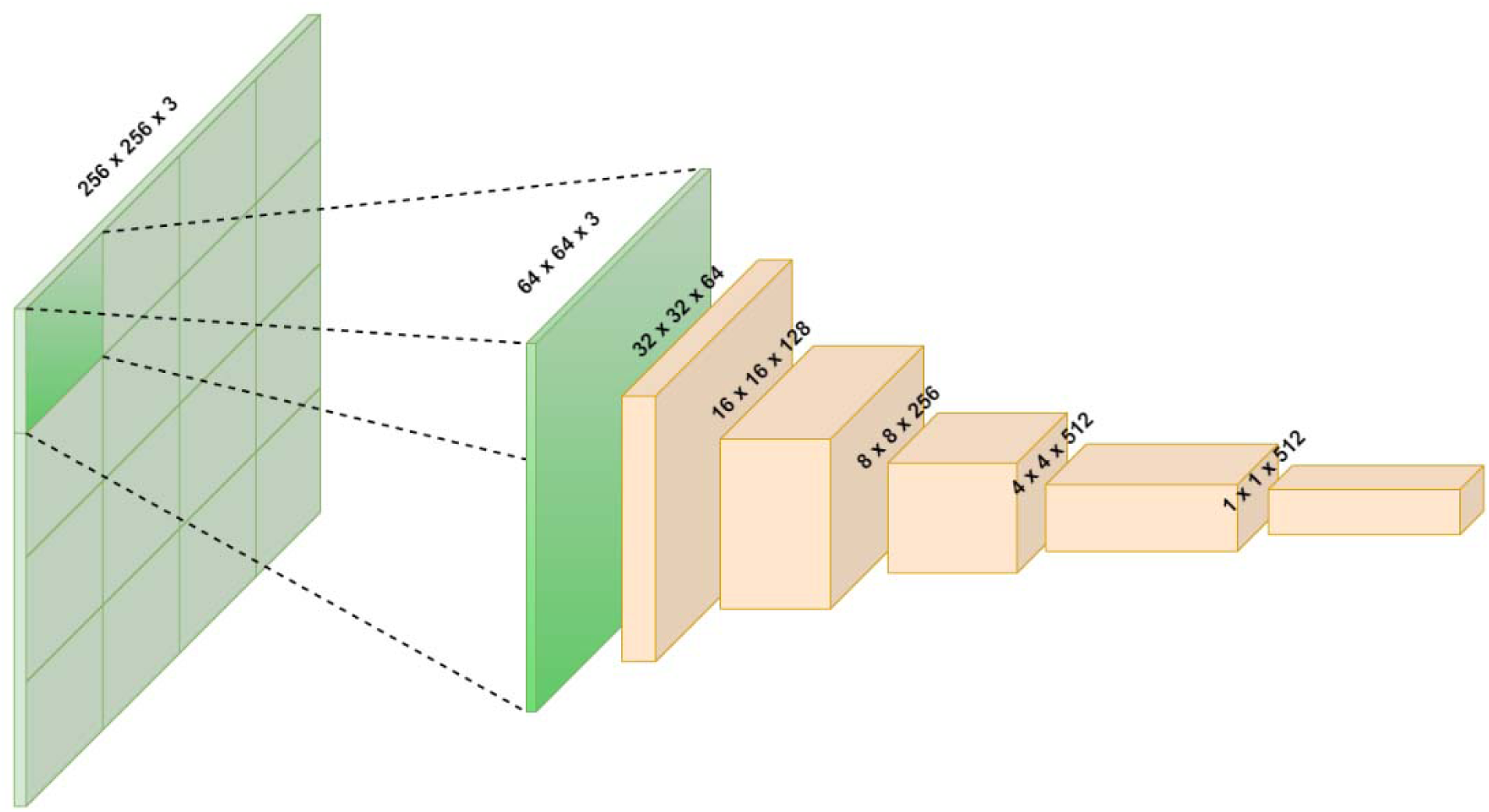
Patch Discriminator Architecture

## Notes

### Competing Interest Statement

The authors have declared no competing interest.

### Summary of Updates

We have updated with more results in data validation and quantification.

